# Situation of antibiotic resistance in Bangladesh and its association with resistance genes for horizontal transfer

**DOI:** 10.1101/2020.04.06.027391

**Authors:** Kazi Sarjana Safain, Golam Sarower Bhuyan, Sadia Tasnim, Saad Hassan Hasib, Rosy Sultana, Mohammad Sazzadul Islam, Mohammad Al Mahmud-Un-Nabi, Suprovath Kumar Sarker, Farjana Akther Noor, Asifuzzaman Rahat, Md Abdul Majid Bhuiyan, Md Tarikul Islam, Farhana Manzoor, Sajid Anwar, Daniel Leung, Syed Saleheen Qadri, Firdausi Qadri, Kaiissar Mannoor

**Affiliations:** Infectious Diseases Laboratory, institute for developing Science and Health initiatives, Mohakhali, Dhaka, Bangladesh; Genetics and Genomics Laboratory, institute for developing Science and Health initiatives, Mohakhali, Dhaka, Bangladesh; Department of Mathematics and Natural Sciences, BRAC University, Mohakhali, Dhaka, Bangladesh; Department of Immunology, Bangladesh University of Health Sciences, Mirpur, Dhaka, Bangladesh; BIHSH and UHC Component Institution, Bangladesh Institute of Health Sciences General Hospital, Mirpur, Dhaka, Bangladesh; Department of Enteric and Respiratory Infectious Diseases, Infectious Diseases Division, International Centre for Diarrhoeal Disease Research, Bangladesh, Mohakhali, Dhaka, Bangladesh; Department of Biochemistry and Molecular Biology, Jagannath University, Dhaka, Bangladesh; Division of Infectious Diseases, The University of Utah, Salt Lake City, United States; Department of Genetic Engineering and Biotechnology, University of Dhaka, Dhaka, Bangladesh; Department of Biochemistry and Molecular Biology, University of Dhaka, Dhaka, Bangladesh

## Abstract

The study investigated the spectrum of antibiotic resistance and the associated genes for aminoglycoside, macrolide and ESBL class of antibiotics using clinical isolates. A total of 430 preserved bacterial strains (*Acinetobacter baumannii*, n= 20; *Pseudomonas aeruginosa*, n= 26; *Klebsiella pneumoniae*, n= 42; *E.coli*, n= 85; *Staphylococcus aureus*, n= 84; *Salmonella* Typhi, n= 82; *Enterococcus* spp., n= 27; *Streptococcus pneumoniae*, n= 36 and CNS, n= 28) were examined. The strains were isolated from patients admitted to various tertiary hospitals of Dhaka city between 2015 and 2019 with either acute respiratory infections, wound infections, typhoid fever or diarrhea. The isolates were reconfirmed by appropriate microbiological and biochemical methods. Antimicrobial susceptibility tests were done using Kirby–Bauer disk diffusion approach. PCR amplification using resistance gene-specific primers for aminoglycoside, macrolide and ESBL class of antibiotics was done and the amplified products were confirmed by Sanger sequencing. Of the total isolates, 53% came out as MDR with 96.6% of *E. coli* and 90% of *Staphylococcus aureus*. There was a year-wise gradual increase of MDR isolates from 2015-2018 and by 2019 the increase in MDR isolates became almost 2-fold compared to 2015. Among the five ESBL genes investigated, CTXM-1 came out as the most prevalent (63%) followed by NDM-1 (22%) and *E. coli* isolates were the predominant reservoir of these genes. ErmB (55%) was the most frequently detected macrolide resistance gene, whereas aac(6)-Ib (35.44%) was the most prevalent aminoglycoside resistance gene and these genes were most prevalent in *E. coli* and *P. aeruginosa* isolates, respectively. CTXM-1 and ErmB (16.66%) were the most frequent partners of coexistence followed by CTXM-1 and aac(3)-II.

## Introduction

Antimicrobial resistance (AMR) is becoming a very challenging problem worldwide. Solving the AMR problem should be the priority of global efforts. Nosocomial infections have been recognized as the hub for thriving multidrug resistant (MDR) pathogens. It is estimated that around 8.7% of the hospitalized patients worldwide develop nosocomial infections which is the leading cause of surgical failure, transplant rejection, treatment failure, increased costs and even deaths [1]. Numerous reports suggest that absolute numbers of infections due to resistant microbes are increasing globally [2-4]. It is conservatively estimated that at least 2 million illnesses and 23,000 deaths had been caused by antibiotic resistant organisms per year in the USA [5]. The present trend predicts that infections by resistant bacterial pathogens may cause up to 10 million deaths/year – more than any other causes – by 2050 and like the most global issues, the problem is inequitably distributed, with approximately 90% of the predicted deaths are estimated to happen in Asia and Africa [6]. Additionally, treatment of the patients infected with resistant pathogens is associated with higher costs, requiring additional investigations and longer hospitalization [7]. The overall crude economic burden of antibiotic resistance was estimated to be at least €1.5 billion in 2007 in Europe and $55 billion in 2000 in the USA, including patients and hospital costs [8]. Without appropriate action, by 2050 the global economy may lose more than USD 6 trillion annually because of AMR, which is nearly 4% of Gross Domestic Product (GDP) [7].

Studies show that low-income countries like Bangladesh are more affected by AMR because of the widespread misuse of antibiotics, non-human antibiotic use, poor quality of drugs, inadequate surveillance and factors associated with individual and national poverty indicators like poor healthcare standards, malnutrition, chronic and repeated infections, unaffordability of more effective and costly drugs [9, 10]. Research regarding antibiotic resistance epidemiology may ultimately guide to interventions for AMR.

Regional surveillance programs have tracked antibiotic resistance in several countries, but still many gaps exist in some middle and low income countries that prohibit comprehensive monitoring and analysis of the prevalence and trends of resistance worldwide to appropriately guide interventions [11]. Thus, a well-functioning national AMR surveillance system is vital in planning and implementing the national AMR strategy. Studies on antimicrobial resistance encompassing different aspects on a national scale are critical because they provide information on the extent of established resistance rates, as well as emerging patterns of resistance. Understanding how resistance is changing over time is important for (1) establishing prescribing guidelines, (2) determining investment in new therapies, and (3) improving the targeting of campaigns to reduce antimicrobial resistance. It also provides a baseline for future analysis and comparison with other countries [12]. Studying resistance genes that cause antibiotic resistance and the plasmids that transfer such resistance genes, and other mechanisms that lead to antibiotic resistance will help to generate new ideas, which in turn may lead to control the spread of resistance by targeting the resistance genes.

Previous studies on antibiotic resistance in Bangladesh have largely focused on Gram-negative bacteria [13-15], and a limited number of antibiotic resistance genes [16-18]. The present study was undertaken to investigate the contemporary antibiotic resistance patterns in Bangladesh, including that of Gram-positive bacteria, which have emerged as global threats in serious infections like pneumonia, wound infections and diabetic ulcer [19-21]. So, the study of antibiotic resistance to Gram-positive clinical isolates is by no means less important. In addition to antibiotic resistance patterns of Gram-positive and Gram-negative bacteria, the study also investigated the genetic basis of resistance by analyzing previously reported resistance genes and a set of other genes that had not been investigated in previous studies in Bangladesh, such as aac(6)-Ib, aac(3)-II, aph(3)-VI, ant(3)-I, and ErmF genes. A comparative analysis of co-presence of resistance genes was also done to know the importance of the presence of multiple genes in the horizontal spread of antibiotic resistance for Gram-negative and Gram-positive bacteria. To know how alarmingly the resistance was increasing with the passage of time, the kinetics of the rate of increase of multidrug resistant (MDR) pathogens was determined between 2015 and 2019.

## Materials and Method

The present prospective study was carried out in the institute for developing Science and Health initiatives (ideSHi), Dhaka, Bangladesh.

### Bacterial isolates

The study analysed a total of 430 preserved bacterial isolates. These isolates were collected from hospitalized patients with acute respiratory infections (nasopharyngeal swab), wound infections (wound swab), typhoid fever (blood samples) and diarrhoeal diseases (stool samples) between 2015 and 2019. The strains were preserved at −70°C as glycerol stock. Bacterial identification and reconfirmation were performed by routine conventional microbial cultures and biochemical tests using standard recommended techniques [22].

### Antimicrobial susceptibility testing

Profiling of antimicrobial susceptibility/resistance was performed by the modified Kirby-Bauer disc diffusion method and the bacterial strains were identified as either sensitive or resistant to an antibiotic based on the diameter of inhibition zone interpretative chart, published in the Clinical and Laboratory Standard Institute (CLSI) guidelines, 2016. MDR strains were identified as resistant to three or more antimicrobial classes [23].

The cartridges of antimicrobial disks for Nalidixic acid, Cotrimoxazole, Amoxiclav, Ciprofloxacin, Azithromycin, Ampicillin, Cefixime, Gentamycin, Chloramphenicol, Meropenem, Ceftriaxone, Imipenem, Piperacillin-tazobactam, Norfloxacin, Tobramycin, Amikacin, Netilmicin, Carbenicillin, Tetracycline, Erythromycin, Levofloxacin, Doxycycline, Streptomycin, Trimethoprim-sulfamethoxazole, Polymyxin B, Rifampicin, Cefoxitin, Penicillin, Vancomycin, Linezolid, and Clindamycin were obtained from Oxoid (Hampshire, UK). The cartridges were stored between 4°C and 20°C and allowed to come to room temperature prior to use. After inoculation with the isolates and placement of the disks, Mueller-Hinton agar plates were incubated at 37°C for 24 hours and the zones of inhibition were measured.

### Determination of MAR index

Multiple antibiotic resistance index (MAR) is broadly helpful in analyzing potential risk, and is used to determine the severity of antibiotic resistance [24]. In this study, antibiotics were used according to CLSI guideline for a particular species. MAR value was defined as a/b, where a represents the number of antibiotics to which the isolate was resistant, and b represents the number of antibiotics to which the isolate was exposed.

### PCR amplifications and sequencing

Bacterial DNA was extracted from the isolated bacterial colonies by boiling method [25]. All the isolates were examined for the presence of resistance genes for different classes of antibiotics, namely aminoglycoside, macrolide and β-lactam. The gene sequences were retrieved from the nucleotide database of National Centre for Biotechnology Information (NCBI). The retrieved FASTA sequences were then used to design gene-specific primers using NCBI Primer-BLAST. The following sets of primers were used for each resistance gene: (1) *aac(3)-II*: F1: 5′-ATATCGCGATGCATACGCGG-3′, R1: 5′-GACGGCCTCTAACCGGAAGG-3′; (2) *aac(6)-Ib*: F2: 5′-TTGCGATGCTCTATGAGTGGCTA-3′, R2: 5′-R1CTCGAATGCCTGGCGTGTTT-3′; (3) *aac(6)-II*: F3: 5′-CGACCATTTCATGTCC-3′, R3: 5′-GAAGGCTTGTCGTGTTT-3′; (4) *ant(3)-I*: F4: 5′-CATCATGAGGGAAGCGGTG-3′, R4: 5′-GACTACCTTGGTGATCTCG-3′; (5)*aph(3)-VI*:F5:5′-ATGGAATTGCCCAATATTATT-3′,R5:5′-TCAATTCAATTCATCAAGTTT-3′; (6) *armA*: F6: 5′-CCGAAATGACAGTTCCTATC-3′, R6: 5′-GAAAATGAGTGCCTTGGAGG-3′; (7) *rmtB*: F7: 5′-ATGAACATCAACGATGCCCTC-3′, R7:5′-CCTTCTGATTGGCTTATCCA-3′;(8)*ErmA*:F8:5′-CTTCGATAGTTTATTAATATTAGT-3′, R8: 5′-TCTAAAAAGCATGTAAAAGAA-3′; (9) *ErmB*: F9: 5′-GGAACATCTGTGGTATGGCG-3′, R9: 5′-CATTTAACGACGAAACTGGC-3′; (10)*ErmC*:F10:5′-CAAACCCGTATTCCACGATT-3′,R10:5′-ATCTTTGAAATCGGCTCAGG-3′; (11) *ErmF*: F11: 5′-TGTTCAAGTTGTCGGTTGTG-3′, R11:5′-CAGGACCTACCTCATAGACA-3′;(12)*KPC*:F12:5′-CTGTCTTGTCTCTCATGGCC-3′, R12: 5′-CCTCGCTGTGCTTGTCATCC-3′; (13) *Oxa48*: F13: 5′-GCGTGGTTAAGGATGAACAC-3′, R13: 5′-CATCAAGTTCAACCCAACCG-3′; (14) *VIM1*: F14: 5′-AGTGGTGAGTATCCGACAG-3′, R14: 5′-ATGAAAGTGCGTGGAGAC-3′; (15)*NDM-1*:F15:5′-GGTTTGGCGATCTGGTTTTC-3′,R15:5′-CGGAATGGCTCATCACGATC-3′;(16)*CTX-M1*:F16:5′-AATCACTGCGCCAGTTCACGCT-3′, R16: 5′-AGCCGCCGACGCTAATACA-3′. Gene-specific PCR amplification using each set of primers was carried out on a T100 ™ thermal cycler (Bio-Rad, USA). The PCR reaction volume was 10 µL containing 1 µL 10X PCR buffer, 0.3 µL 50mM MgCl_2_, 0.2 µL of 10mM dNTPs mixture, 0.5 µL forward and reverse primers, 0.05 µL of Taq polymerase, 5.45 µL Nuclease Free Water and 2 µL of template DNA. The following cycling conditions were used: initial denaturation at 95°C for 5 minutes, 35 cycles of denaturation at 95°C for 30 seconds, annealing at 55°C, cyclic extension at 72°C for 45 seconds, and final extension at 72°C for 6 minutes. The amplified PCR products were visualized on a 1% agarose gel stained with SYBR Safe (Invitrogen, USA) staining. Later, the PCR products were purified using Quick PCR product purification kit (Invitrogen, USA) following the manufacturer’s guidelines.

### Sanger DNA sequencing

The PCR-amplified DNA was sequenced using ABI PRISM-310 software version 3.1.0 (Applied Biosystems). To prepare cycle sequencing reaction mixtures, 1 ng of purified DNA was added to a reaction solution containing 0.50 µL Big dye terminator sequencing mix and 2.0 µL of 5X PCR sequencing buffer with 0.2 µL primer solution. Then, the PCR strip was placed in the Mastercycler^®^ gradient Thermal Cycler (Cat. No. 4095-0015, USA Scientific) and subjected to the following thermal cycling profile: pre-denaturation at 95°C for 10 minutes; 25 cycles of denaturation at 95°C for 10 seconds, annealing at 55°C for 5 seconds, cyclic extension at 72°C for 4 minutes; and a final extension at 72°C for 6 minutes.

### Data analysis

Sequencing data were analysed by Chromas Lite 2.4 software to identify the target sequence by alignment with reference sequence. The obtained sequence was subjected to further analysis using Basic Local Alignment Search Tool (BLAST) for finding sequence similarity with sequences already reported in online databases. Graphs were generated using GraphPad Prism 7.00 software.

## Results

### Selection of clinical isolates

The study analyzed a total of 430 preserved (−80°C) clinical bacterial isolates from patients with acute respiratory infections (n=40), wound infections (n=239), typhoid (n=82) and diarrhea (n=69). Among the enlisted 12 global priority pathogens (GPP) according to World Health Organization (WHO), the present study worked on 8 bacterial species from GPP list including *Acinetobacter baumannii* (n=20), *Pseudomonas aeruginosa* (n=26), *Klebsiella pneumoniae* (n=42) and *Escherichia coli* (n=85) as the Priority 1 (Critical) pathogens, followed by *Staphylococcus aureus* (n=84*), Salmonella* Typhi (n=82) and *Enterococcus* spp. (n=27) as the Priority 2 (High) pathogens. Among the Priority 3 (Medium) pathogens list, *Streptococcus pneumoniae* (n=36) isolates were analyzed. Additionally, CNS (n=28) which is not enlisted in any of the priority list, was also investigated. These clinical specimens were collected from patients admitted to various tertiary hospitals of Dhaka city between 2015 and 2019 (Table 1).

**Table 1.** Information of clinical isolates analyzed.

| Clinical isolates | *Pseudomonas aeruginosa* | *Klebsiella pneumoniae* | *Acinetobacter baumannii* | *Escherichia coli* | *Salmonella* Typhi | *Enterococcus* spp. | *Staphylococcus aureus* | *Streptococcus pneumoniae* | CNS |
| --- | --- | --- | --- | --- | --- | --- | --- | --- | --- |
| Typhoid (n= 82) |  |  |  |  | 82 |  |  |  |  |
| Diarrhea (n=69) |  |  |  | 69 |  |  |  |  |  |
| Wound infection (n= 239) | 26 | 30 | 20 | 12 |  | 17 | 84 | 12 | 28 |
| Acute respiratory |  | 12 |  | 4 |  |  |  | 24 |  |
| infection<br>(n= 40) |  |  |  |  |  |  |  |  |  |
| Total (n= 430) | Total (n= 26) | Total (n= 42) | Total (n= 20) | Total (n= 85) | Total (n= 82) | Total (n= 27) | Total (n= 84) | Total (n= 36) | Total (n= 28) |

<sup>a</sup>CNS: Coagulase-negative *Staphylococcus*

### Antibiotic susceptibility/resistance patterns for the isolated bacterial strains

Next, we wanted to see the antibiotic resistance pattern of the isolated Gram-negative (Figs 1A-1E) and Gram-positive (Figs 1F-1J) bacterial isolates. To interpret the data, the median value of the resistance rate of each of the isolates was calculated and the data were analyzed. In Fig 1(A), we see that *S*. Typhi, which is the major causative agent for typhoid, was found to be 89% resistant to nalidixic acid. However, 10 out of 19 antibiotics showed complete sensitivity to *S*. Typhi. Moreover, *P. aeruginosa* and *K. pneumoniae* isolates exhibited an elevated resistance to almost all the antibiotics tested. Overall, *P. aeruginosa* showed the highest resistance to majority of the antibiotics investigated. For *P. aeruginosa*, 15 antibiotics were tested and all of these antibiotics exhibited resistance at different degrees and 7 of these antibiotics including imipenem (45%), ciprofloxacin (59%), tobramycin (78%), gentamycin (67%), amikacin (45%), netilmicin (45%) and carbenicillin (48%) showed higher resistance rates than the median (41%) (Fig 1B). Next, 7 out of 18 antibiotics tested (ceftriaxone, ciprofloxacin, azithromycin, cefixime, tobramycin, ampicillin and gentamycin) against *K. pneumoniae* crossed the median (22%) with resistance rates of 40%, 47%, 58%, 66%, 42%, 33% and 42%, respectively (Fig 1C). For Gram-positive bacteria, although there were antibiotics which exhibited enough efficacy against *S. aureus* such as vancomycin (4% resistance) and chloramphenicol (3% resistance), still 8 out of 16 antibiotics crossed the median (24%) and 4 antibiotics, namely cefixime, cefoxitin, erythromycin, and penicillin had more than 50% resistance. In addition, 84% *S. aureus* isolates were found to be MRSA (Methicillin Resistant *S. aureus*) as these *S. aureus* strains were resistant to cefoxitin (Fig 1F). For CNS, 7 out of 18 antibiotics including ciprofloxacin, rifampicin, cefoxitin, erythromycin, penicillin, levofloxacin and clindamycin crossed the median (33%) with resistance rates of 38%, 38%, 41%, 58%, 66%, 38% and 38%, respectively (Fig 1G). Overall, the data reveal that although there were 1 or more antibiotics which showed complete sensitivity against *S*. Typhi, *E. coli, A. baumannii, Enterococcus* spp. and *S. pneumoniae*, majority of the tested antibiotics showed resistance from moderate to extremely higher degrees against these organisms. On the contrary, *P. aeruginosa, K. pneumoniae, S. aureus*, and CNS isolates showed resistance to all of the tested antibiotics with different degrees and even many of these bacterial isolates exhibited more than 50% resistance, which may indicate an alarming and deteriorating situation for antibiotic use. Colistin resistance was detected in 10% *K. pneumoniae* and 11% *P. aeruginosa*. Finally, like Gram-negative bacteria, Gram-positive bacteria had equally raised resistance against different antimicrobials.

**Fig 1.**
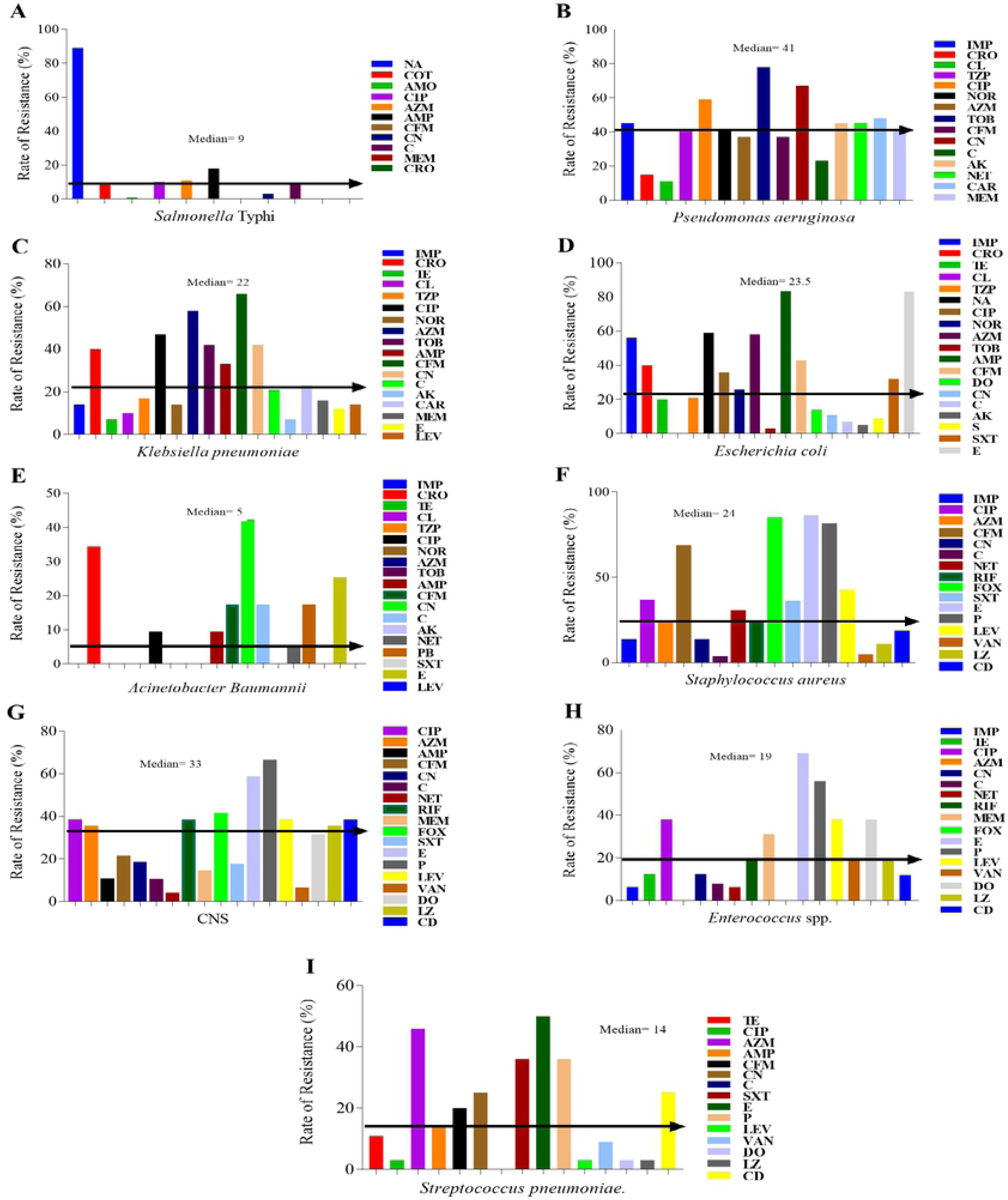
Rates of resistance of the isolated organisms to different antibiotics. Kirby Bauer disk diffusion method was applied to determine resistance/sensitivity rate of each isolate. A. *Salmonella* Typhi, B. *Pseudomonas aeruginosa*, C. *Klebsiella pneumoniae*, D. *E. coli*, E. *Acinetobacter baumannii*, F. *Staphylococcus aureus*, G. CNS, H. *Enterococcus* spp., I. *Streptococcus pneumoniae*. Abbreviation expansion for antibiotics used: NA= Nalidixic acid, COT= Cotrimoxazole, AMO= Amoxiclav, CIP= Ciprofloxacin, AZM= Azithromycin, AMP= Ampicillin, CFM= Cefixime, CN= Gentamycin, C= Chloramphenicol, MEM= Meropenem, CRO= Ceftriaxone, IMP= Imipenem, CL= Colistin, TZP= Piperacillin-tazobactam, NOR= Norfloxacin, TOB= Tobramycin, AK= Amikacin, NET= Netilmicin, CAR= Carbenicillin, TE= Tetracycline, E= Erythromycin, LEV= Levofloxacin, DO= Doxycycline, S= Streptomycin, SXT= Trimethoprim-sulfamethoxazole, PB= Polymyxin B, RIF= Rifampicin, FOX= Cefoxitin, P= Penicillin, VAN= Vancomycin, LZ= Linezolid, CD= Clindamycin.

### Antibiotic resistance profile and MAR index

Table 2 shows the antibiotic resistance profile and Multiple Antibiotic Resistance (MAR) index of the indicated isolates. The proportion of the isolates with MAR index >0.2 was 53.73%; while those with MAR index ≤ 0.2 was 43.25% and the findings indicate that more than 50% of the isolates were likely to be from high risk sources. A large proportion of *P. aeruginosa* (80.77%), *S. aureus* (78.57%) and *K. pneumoniae* (76.2%) isolates exhibited MAR index value of >0.2, whereas the highest mean value of MAR index (0.562±18) was observed for *P. aeruginosa*.

**Table 2.** Distribution of Multiple Antibiotic Resistance (MAR) indices within the identified organisms.

| MAR Index (Range) | *S. Typhi*,<br>n (%) | *P. aeruginosa*<br>n (%) | *K. pneumoniae*<br>n (%) | *E. coli*<br>n (%) | *A. baumannii*<br>n (%) | CNS<br>n (%) | *E. spp*<br>n (%) | *S. aureus*<br>n (%) | *S. pneumoniae*<br>n (%) | Total<br>n (%) |
| --- | --- | --- | --- | --- | --- | --- | --- | --- | --- | --- |
| <0.2 | 52 (63.42) | 5 (19.23) | 10 (23.8) | 56 (65.89) | 9 (45) | 9 (32.14) | 13 (48.15) | 14 (17.86) | 17 (47.22) | 186 (43.25) |
| 0.2 | 0 | 0 | 0 | 0 | 0 | 3 (10.75) | 0 | 3 (3.57) | 7 (19.44) | 13 (3.02) |
| >0.2 | 30 (36.58) | 21 (80.77) | 32 (76.2) | 29 (34.11) | 11 (55) | 16 (57.11) | 14 (51.85) | 66 (78.57) | 12 (33.34) | 231 (53.73) |
| Mean±SD | 0.34±0.108 | 0.562±0.18 | 0.38±0.15 | 0.32±0.133 | 0.312±0.089 | 0.45±0.11 | 0.38±0.14 | 0.4±0.11 | 0.34±0.059 | |
| Total | 82 | 26 | 42 | 85 | 20 | 28 | 27 | 84 | 36 | 430 |

### Frequencies of MDR isolates

To evaluate the extent of multidrug resistance, the percentage of resistance for each bacterial species was assessed. Approximately, 53% isolates were identified as MDR. Fig 2A shows that *E. coli* (96.6%) was found to be the most prevalent MDR organism, followed by *S. aureus* (90%), *P. aeruginosa* (83.4%), *K. pneumoniae* (80.69%), CNS (75.5%), *S. aureus* (75%), *A. baumannii* (66.67%), *Enterococcus* spp. (60.42%), *S. pneumoniae* (55.62%) and *S*. Typhi (34.7%). Next, in Fig 2(B), we see that resistance with more than 6 antibiotics was found to be 68% for *S. aureus*, followed by 62.5% for *P. aeruginosa* and 5% for CNS. The resistance rates for 2, 3, 4, 5, 6, and >6 antibiotics were 3.4%, 10.22%, 28.4%, 22.72%, 11.4% and 23.86%, respectively among the *E. coli* isolates. Among *P. aeruginosa* and *K. pneumoniae*, only 2*%* isolates exhibited 100% sensitivity. The resistance rates for 1, 2, 3, 4, 5, 6, and >6 antibiotics were 6.3%, 8.3%, 0%, 8.3%, 8.3%, 4.3% and 62.5%, respectively for the *P. aeruginosa* isolates, whereas the corresponding resistance rates for *K. pneumoniae* were 10%, 7.31%, 2.41%, 17%, 9.75%, 12.53% and 39%. On the other hand, 1%, 2%, 1.6%, 0.4%, 5%, 21% and 68% *S. aureus* isolates were found to be resistant to 1, 2, 3, 4, 5, 6, and >6 antibiotics, respectively. So, the data indicates that antibiotic resistance levels have reached alarmingly high levels for *S. aureus, P. aeruginosa, K. pneumoniae* and *E. coli*, although *S*. Typhi and *S. pneumoniae* isolates had shown susceptibility to most of the antibiotics tested and none of the isolates was resistant to >6 antibiotics.

**Fig 2.**
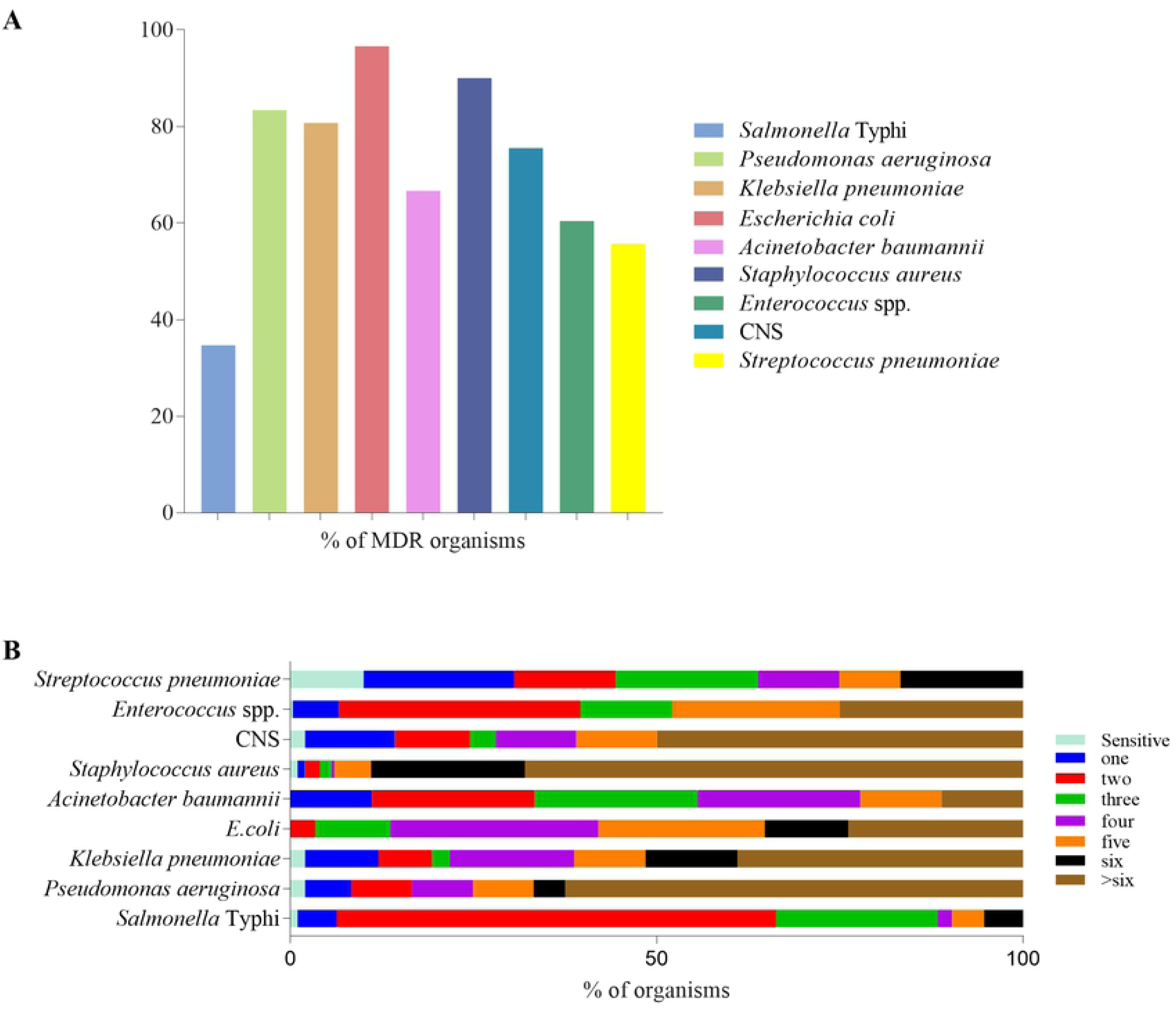
Resistance frequency of the indicated clinical isolates. (A) MDR frequencies of the indicated organisms. (B) Proportion of the mentioned bacterial isolates which were resistant to the indicated numbers of drugs. By % MDR in Fig A, we mean % isolates showing resistance to more than 3 antibiotic classes among total isolates of a bacterial species and each color here indicates a bacterial species. For Fig B, each color indicates the number of antibiotics to which a bacterial species was resistant.

### Resistance gene profiles among the tested isolates

To observe the resistance gene spectrum among different bacterial isolates, the percentages of antibiotic resistance genes were assessed. Fig 3 shows that *E. coli* strains had harbored all of the resistance genes analyzed except aph(3)-VI. *S. aureus, K. pneumoniae* and *P. aeruginosa* were detected as the next prevalent organisms which had higher percentages of resistance genes, in particular CTXM-1, ErmB, aac(6)-Ib and aac(3)-II. The most frequently identified ESBL gene was CTXM-1 (Amber class-A type) and it was detected in almost every species of bacteria with the highest frequencies found in *S. aureus* (36%), followed by *K. pneumoniae* (30%) and *E. coli* (28%). However, none of the *S*. Typhi and CNS strains showed to harbor this gene. NDM-1, which is a metallo-β-lactamase of Amber class-B was also shown to be commonly detected among ESBL genes. The predominant bacterial species which harbored this gene was found to be *K. pneumoniae* (19%). Among the Amber class-D type ESBL genes, OXA-48 was predominantly present in 14.75% *S. aureus* strains. The commonly detected macrolide resistance gene, ErmB was detected in 24.50% *E. coli*, 19% *K. pneumoniae* and 16% *P. aeruginosa*. In addition, 30% *P. aeruginosa* and 11% CNS were found to possess aac(6)-Ib gene, which is the frequently detected aminoglycoside resistance gene. *S. pneumoniae, S*. Typhi and *Enterococcus* spp. were found to harbor only 2 of the investigated resistance genes, with ErmB being the commonly harbored resistance gene.

**Fig 3.**
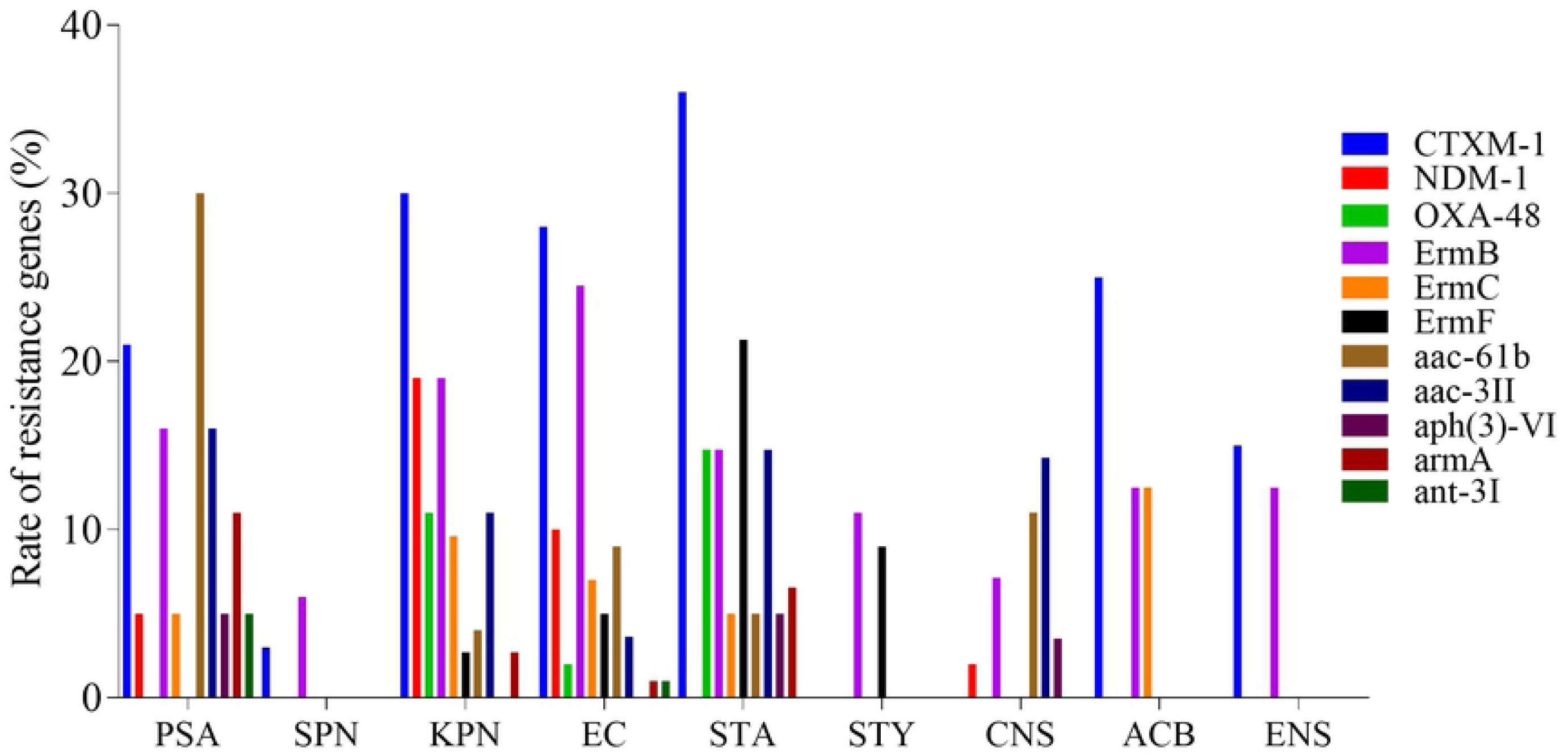
Distribution of CTXM-1, NDM-1, OXA-48, ErmB, ErmF, ErmC, aac(6)-Ib, aac(3)-II, aph(3)-VI, armA and ant-3I genes among the indicated isolates. In the Figure, PSA= *Pseudomonas aeruginosa*, SPN= *Streptococcus pneumoniae*, KPN*= Klebsiella pneumoniae*, EC*= Escherichia coli*, STA=*Staphylococcus aureus*, STY*= Salmonella* Typhi, ACB= *Acinetobacter baumannii*, ENS= *Enterococcus* spp.

### Phenotypic discrepancies

The antibiotic-resistant determinants associated with resistance were detected by PCR and subsequently confirmed by Sanger sequencing. Of the 430 isolates tested, 115 (27%) isolates did not harbor any of the genes tested even though they were phenotypically resistant by disk diffusion (the green bar of Fig 4). On the contrary, 10 (2.33%) of the total isolates encompassing all bacterial species were observed to be phenotypically inconsistent although these isolates harbored resistance genes. The rest of the isolates (n=305) had matching consistency in terms of genotypes and phenotypes.

**Fig 4.**
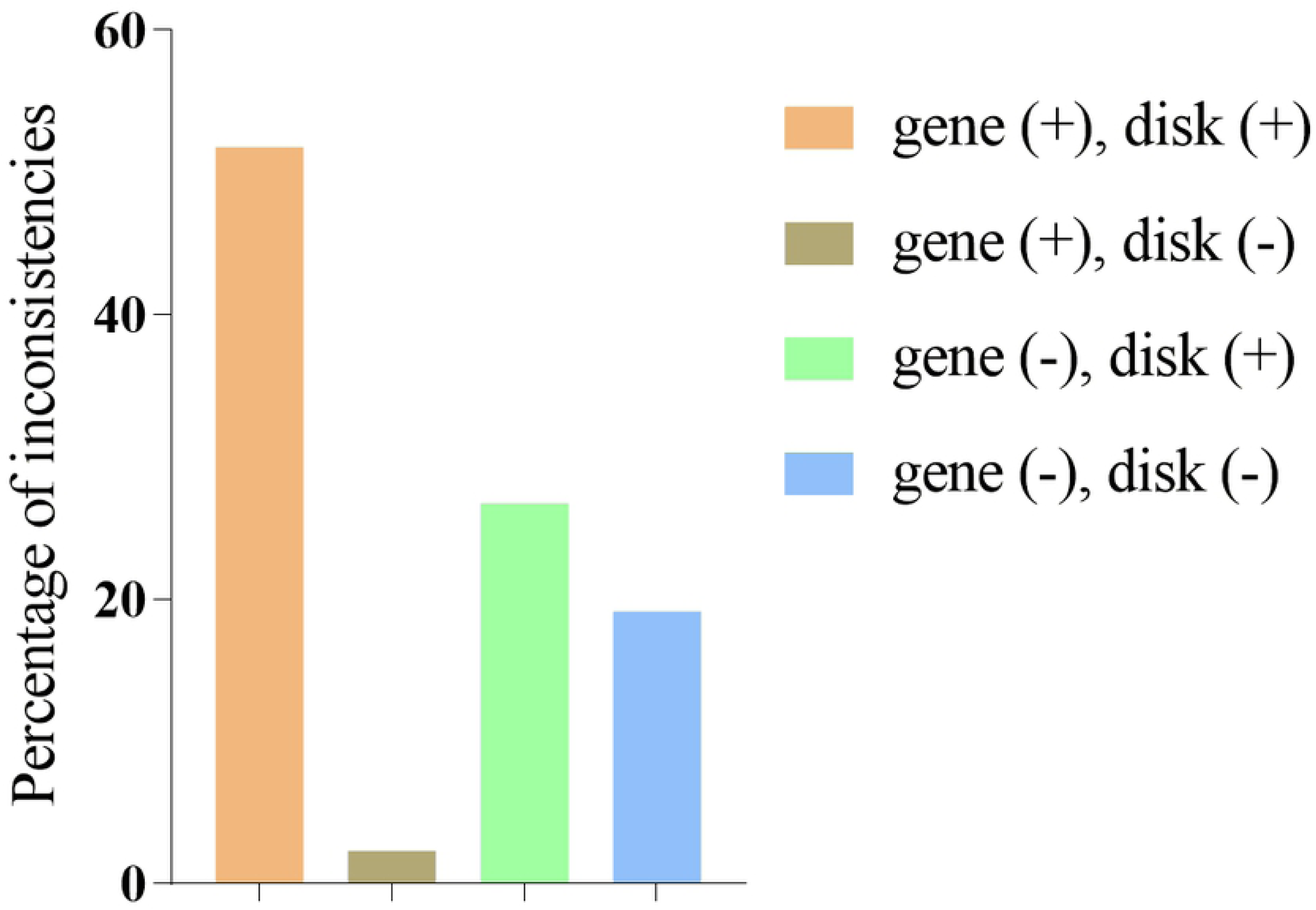
Existing inconsistencies among the isolated organisms. The orange bar indicates the organisms having their resistance genes detected and these organisms gave positive results on disk diffusion analysis, whereas the brown bar indicates those organisms having their resistance genes detected but the disk diffusion results were negative. On the other hand, the green bar indicates organisms for which no resistance genes were detected but they showed resistance on disk diffusion analysis, while the blue bar indicates those isolates which gave negative results in terms of both gene detection and antibiogram results.

### Comparison of the year-wise rate of resistance genes and MDR isolates

To observe the antibiotic resistance trend of the pathogenic organisms enlisted in the WHO priority list, the year-wise percentages of MDR organisms along with the resistance genes were compared between 2015 and 2019. Fig 5A shows that the percentages of organisms harboring CTXM-1, ErmB, aac(6)-Ib, NDM-1, and aac(3)-II resistance genes were 14, 17, 22, 3, and 3 for 2015 specimens, whereas the presence of these resistance genes increased to 44%, 35%, 28%, 16%, and 29%, respectively for specimens of 2019. On the other hand, although the resistance genes, namely OXA-48, ErmC, ErmF, aph(3)-VI, and armA were not detected in specimens of 2015, they appeared in 10%, 2%, 10%, 5%, and 3% of the tested organisms of 2019. Next, Fig 5B shows that MDR organisms were gradually increasing with the passage of time. There were 33% (n=30) MDR organisms in 2015 and they increased gradually in 2016, 2017, and 2018 and became 62% (n=67) in 2019. That is, the proportion of MDR organisms has become almost double in 2019 compared to the 2015. Overall, the data reveal that rates of MDR organism along with resistance genes are in an alarmingly increasing trend with the passage of time.

**Fig 5.**
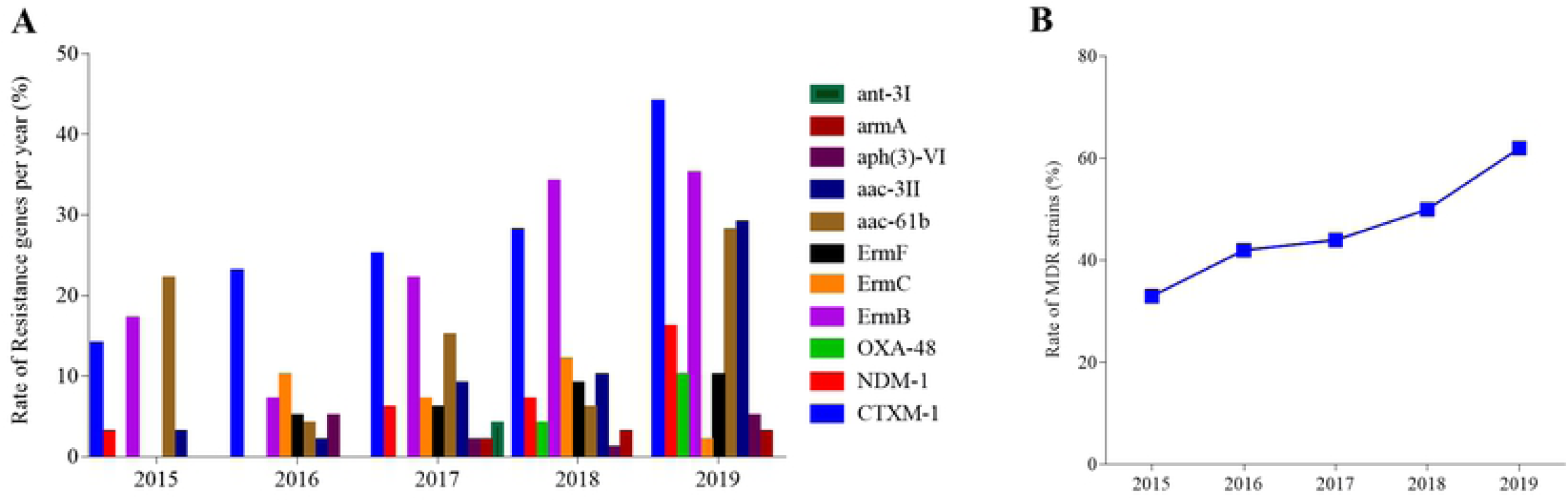
Kinetics of the spread of MDR organisms along with antibiotic resistance genes upon passage of time. (A) Comparison of the percentages of the organisms carrying antibiotic resistance genes over five years, (B) Comparison of the rate of sums of the MDR organisms (resistant to more than 3 antibiotic classes) that included PSA, SPN, KPN, EC, STA, STY, CNS, ACB, ENS isolates between 2015 and 2019. For both Fig A and Fig B, the number of samples analyzed were 92, 82, 63, 85 and 108 for 2015, 2016, 2017, 2018 and 2019, respectively. Each color in Fig A indicates an antibiotic resistance gene.

### Co-existence of multiple antibiotic resistance genes in the tested isolates

Next, we wanted to investigate co-presence of resistance genes. The results of co-presence of resistance genes are displayed in Table 3. Out of 430 strains, a total of 102 strains revealed to possess co-presence of resistance genes. *E. coli* (n=40), *P. aeruginosa* (n=26) and *S. aureus* (n=17) strains were found to be the predominant organisms in harboring the co-resistance genes. In the 85 *E. coli* isolates analyzed, co-existence of (a) CTXM-1 and ErmB (n=10) was identified as the most prevalent combination, followed by (b) CTXM-1 and aac(3)-II (n=5), (c) ErmB and ErmC (n=4), (d) CTXM-1 and ErmC (n=3), CTXM-1 and ErmF (n=3), ErmB and ErmF (n=3), (e) CTXM-1 and aac(6)-Ib (n=2) and NDM-1and aac(3)-II (n=2). Alarming enough to know that an *E. coli* isolate showed to possess a set of five genes, namely ErmB, ErmF, aac(6)-Ib, ant(3)-I and CTXM-1. On the other hand, all 26 (100%) *P. aeruginosa* isolates that had been analyzed showed co-presence of resistance genes in different combinations. A *P. aeruginosa* isolate was seen to contain multiple ESBL genes (CTXM-1, NDM-1 and OXA-48), with the addition of aac(6)-Ib, an aminoglycoside resistance gene. Overall, the data demonstrate that *P. aeruginosa* has a common phenomenon (100%) of harboring multiple resistance genes, whereas 47% of *E. coli*, 32% of *K. pneumoniae*, and 20% of *S. aureus* isolates harbored multiple genes. The rest of the bacterial species had relatively low presence of multiple genes (Table 3). *S*. Typhi, *S. pneumoniae*, and *Enterococcus* spp. did not have co-presence of any of the tested genes (data not shown). Overall, the data reveal that huge number of Gram-negative *E. coli, P. aeruginosa* and *K. pneumoniae* isolates had co-presence of resistance genes. On the other hand, only *S. aureus* isolates among the Gram-positive bacteria had significant co-presence of resistance genes.

**Table 3.** Detection of the presence of co-resistance genes in the tested organisms. To determine the co-presence of resistance genes, a total of 430 specimens were analyzed by PCR and Sanger sequencing followed by bioinformatics using NCBI blast. Isolates analyzed: *E. coli (n=85), P. aeruginosa* (n=26), *K. pneumoniae* (n=42), *S. aureus* (n=84), *A. baumannii* (n=20) and CNS (n=28).

| Co-existence of resistance genes | *E. coli* | *P. aeruginosa* | *K. pneumoniae* | *S. aureus* | *A. baumannii* | CNS | No. of isolates with co-presence of resistance genes |
| --- | --- | --- | --- | --- | --- | --- | --- |
| CTXM-1, ErmB | 10 |  | 4 | 3 |  |  | 17 |
| CTXM-1, ErmC | 3 |  |  | 2 | 1 |  | 6 |
| CTXM-1, ErmF | 3 | 4 |  |  |  |  | 7 |
| ErmB, ErmC | 4 | 2 |  |  |  |  | 6 |
| ErmB, ErmF | 3 |  |  | 3 | 1 |  | 7 |
| ErmB, ErmC, OXA-48 | 1 |  |  | 1 |  |  | 2 |
| **ErmB, OXA-48** |  | 2 |  |  |  |  | 2 |
| **CTXM-1, aac(6)-Ib** | 1 | 2 | 4 | 1 |  |  | 8 |
| **CTXM-1, NDM-1, aac(3)-II, ErmB** | 1 |  |  | 1 |  |  | 2 |
| **ErmB, ErmF, aac(6)-Ib** | 1 |  |  |  |  |  | 1 |
| **ErmB, aac(3)-II, aph(3)-VI, aac(6)-Ib** |  | 1 |  |  |  | 1 | 2 |
| **ErmB, ErmF, aac(6)-Ib, ant(3)-I, CTXM-1** | 1 |  |  |  |  |  | 1 |
| **ErmB, CTXM-1, aac(3)-II** | 1 | 2 |  |  |  |  | 3 |
| **ErmF, CTXM-1, aac(6)-Ib** |  |  | 1 | 1 |  |  | 2 |
| **Aac(6)-Ib, aac(3)-II** | 2 | 1 | 1 |  |  |  | 4 |
| **Aac(6)-Ib, aac(3)-II, CTXM-1** | 1 |  |  | 1 |  |  | 2 |
| **OXA-48, armA** |  | 1 |  |  |  |  | 1 |
| **OXA-48,**<br>**armA, aac(6)-**<br>**Ib, aac(3)-II** |  |  |  | 1 |  |  | 1 |
| **CTXM-1,**<br>**NDM-1, OXA-**<br>**48, aac(6)-Ib** |  | 1 |  |  |  |  | 1 |
| **CTXM-1,**<br>**NDM-1,**<br>**aac(6)-Ib** |  | 2 | 2 |  |  |  | 4 |
| **NDM-1,**<br>**aac(3)-II** | 1 | 1 |  |  |  |  | 2 |
| **NDM-1,**<br>**aac(6)-Ib** |  |  | 1 |  |  |  | 1 |
| **Aph(3)-VI,**<br>**aac(3)-II,**<br>**CTXM-1** |  |  |  | 2 |  |  | 2 |
| **armA, aac(3)-**<br>**II, CTXM-1** |  |  |  | 1 |  |  | 1 |
| **CTXM-1,**<br>**aac(3)-II** | 5 | 4 |  | 1 |  |  | 10 |
| **OXA-48,**<br>**aac(3)-II** |  | 1 |  | 1 |  |  | 2 |
| **NDM-1,**<br>**aac(3)-II** | 2 | 2 |  |  |  | 1 | 5 |
| **Total** | 40 | 26 | 13 | 17 | 2 | 2 | 102 |

<sup>a</sup>Although *S. Typhi* (n=82), *S. pneumoniae* (36), and *Enterococcus* spp. (n=27) were analyzed but the data are not shown here as co-presence of the tested resistance genes were not detected in these organisms.

## Discussion

The study demonstrates the antibiotic resistance spectrum of the Gram-positive and Gram-negative bacterial isolates from patients with wound infections, acute respiratory infections, diarrhea and typhoid fever. The study also identified the common genes responsible for the resistance of β-lactam, macrolide and aminoglycoside class of antibiotics. Although there are reports regarding antibiotic resistance spectrum for Gram-negative bacteria, no previous work provided antibiotic susceptibilty data on Gram-positive bacteria in Bangladesh. Data on genetic basis of antibitic resistance is also limited in Bangladesh. So, the study was undertaken to address this critical knowledge gap. Major focal points of the study included in vitro antibiotic resistance/sensitivity assay for the preserved strains and determination of the (a) spectrum of multidrug resistance and multiple antibiotic resistance (MAR) indices, (b) antibiotic resistance genes, (c) co-presence of multiple antibiotic resistance genes, and (d) kinetics of the rate of spread of resistance over time. Accurate knowledge on these points would help clinicians to avoid unnecessary use of antibiotics, which in turn would help control of rapid spread of antibiotic resistant bacteria. In addition, the data collected from this report would provide an up-to-date overview of the present situation of antibiotic resistance in Bangladesh and the information in turn will assist in fighting the pathogens more effectively.

The study found that wound infection alone was associated with 7 of the total 12 pathogens from WHO-enlisted global priority pathogens (GPP) list. Moreover, pathogens from wound infections also yielded high rates of ESBL producers such as *E. coli, K. pneumoniae, P. aeruginosa*, and *A. baumannii*. This finding is in line with another study performed in Bangladesh [26]. A large number of wound patients including patients with burns are managed in tertiary care hospitals having causality and surgical departments, and burn units. In spite of technological advances that have been made in surgery and wound management, wound infections have been regarded as the most common nosocomial infections, especially in patients undergoing surgery [27]. Moreover, in Bangladesh empirical use of antiobiotics without confirming the bacterial identification is a common practice and such practices should be strictly regulated.

With exception of *S. pneumaniae, Enterococcus* spp. and *A. baumannii*, most of the isolates irrespective of Gram-negative and Gram-positive bacteria exhibited alarming levels of resistance against majority of the available first line of antibiotics. Furthermore, *S. aureus, K. pneumoniae* and *P. aeruginosa* displayed resistance not only against all the first-line of antibiotics but also against most of the second-line antibiotics tested. These findings are in conformity with the findings of multiple resistance genes in the organisms. For example, although *S. pneumaniae, Enterococcus* spp. and *A. baumannii* had little or no co-presence of resistance genes; *S. aureus, K. pneumoniae* and *P. aeruginosa* had higher levels of co-presence of resistance genes and even 100% of *P. aeruginosa* isolates harbored multiples resistance genes. Such highly resistant organisms may provide a conducive niche for horizonatal transfer of plasmids carrying resistance genes [28], and such a niche may have potential for biofilm formation by the resident bacteria, which in turn may help horizonatal gene transfer [29].

We found that *S*. Typhi isolates were sensitive to several antibiotics, supporting the previous reports [30, 31]. However, although very few of our *S*. Typhi isolates showed resistance to ciprofloxacin or ceftriaxone, WGS analysis of *S*. Typhi isolates by Tanmoy *et al*. showed that appearance of local sublineages of *S*. Typhi with acquisition of new genes, such as qnr and blaCTX-M-15 gene in Bangladesh [32] is making *S*. Typhi treatment complicated. Thus, therapeutic failures in typhoid fever management are not rare in the country [33, 34].

Our study revealed that colistin, the last resort antibiotic, was the most active antimicrobial agent against majority of Gram-negative bacteria, although colistin-resistant pathogens including *K. pneumoniae* and *P. aeruginosa* were detected. In addition to colistin, the antibiotics that were being commonly used previously, such as chloramphenicol and trimethoprim-sulfamethoxazole had relatively higher antimicrobial efficacy, which is in consistence with a single-center study in India [12]. These changes are likely due to the replacement of these drugs as an empiric treatment option for infectious diseases with newer drugs such as aminoglycosides, macrolides and β-lactams. The findings suggest that these drugs can no longer be considered as an empiric treatment option for suspected infections; rather, physicians should rethink to use older drugs again or third-generation cephalosporin. Furthermore, the study showing vancomycin as the strongest antimicrobial against Gram-positive organisms is supported by other studies [35].

Our study revealed CTXM-1 to be the commonest resistance gene (63%), followed by NDM-1 (22%) and OXA-48 (15%) among the ESBL genes. This is in agreement with a 2013-study conducted in Bangladesh which identified CTXM-1 in 50% of the cases [36]. This increase in resistance gene frequencies within a short span of duration (5 years) highlights the contribution of CTXM-1 gene in generation of resistance. *E. coli* was found to be the main reservoir of CTXM-1 gene. CTX-M-mediated genes have also extensively detected in wild birds and poultry [37]; and environmental water [13] in Bangladesh. This provides a vicious cycle of such carbapenemase-producing resistance genes in environment, poultry and human. The finding strengthens the previous report demonstrating CTX-M-producing *E. coli* as a major cause of both community-onset and hospital-acquired infections in Asian countries, including Korea [38], Taiwan [39], China [40], Hong Kong [41], Japan [42], Malaysia [43], and Thailand [44], and the incidence of serious infections due to CTX-M-producing *E. coli* likely will continue to increase [45]. β-lactam antibiotics, especially cefotaxime is very frequently used as first line therapy, which is leading to an alarming escalation of CTX-M-mediated resistance.

After CTXM-1, the resistance gene NDM-1 showed an increasing trend between 2015 and 2019. A recent study showed NDM-1 (55%) as the most prevalent resistance gene in Bangladesh [17]. However, it did not contradict our findings because CTXM-1 was not analyzed in that study. NDM-producing organisms had been reported to be prevalent among *Enterobacteriaceae* in India and Pakistan, even in community-onset infections and is the most dangerous metallo-β-lactamase [46]. This scenario of high NDM-1-producing organism in Bangladesh is of great concern because there are very few new antibiotics in the pharmaceutical pipeline that might work effectively against Gram-negative and none of them will be active against NDM-1 producers [47]. Prior studies detected ESBL genes only in the Gram-negative bacteria. However, our study discovered that these resistance genes existed almost equally in the Gram-positive organisms, also. This shift may result from the horizontal gene transfer which is expected to occur vigorously in organisms causing nosocomical infections.

Also, among the macrolide resistance genes, ErmB (55%) was the most prevalent type. ErmB is mostly associated with azithromycin resistance. In Bangladesh, most diseases such as pneumonia, diarrhea and typhoid are treated with azithromycin, an antibiotic readily available over the counter. The observed frequent detection of ErmB appeared concordant with results from studies reporting ErmB from *S*. Typhi [15], *S. pneumoniae* [16] and *Shigella sonnei* [48]. In a study performed in Bangladesh, Only ErmB was detected among the 7 macrolide resistance genes investigated [15]. Our study could detect the other macrolide resistance genes namely ErmC and ErmF. Moreover, the study detected macrolide resistance genes in *S*. Typhi strains and these findings are of great concern in Bangladesh where both typhoid and azithromycin are common terms to the physicians [15].

The study also found that MDR isolates harboring antibiotic resistance genes increased during the period between 2015 and 2019. CTXM-1, ErmB, aac(6)-Ib, OXA-48, NDM-1, and aac(3)-II had been the major prevalent resistance genes, which were expected to contribute to the alarminly increased levels of spread of MDR organisms. Other studies also confirmed that the relative proportion of bacteria harboring different antibiotic resistance genes (ARG) was broadly increasing over time and since 1970, there was a 15-fold increase of some resistant bacteria by 2008 [49].

Co-presence of multiple resistance genes is a big concern in the spread of antibiotic resistance. Our study found 23.7% strains harboring more than one resistance genes, with an *E. coli* isolate possessing 5 resistance genes. Another study identified more than one carbapenemase gene in 45% of the clinical isolates which had been collected from a hospital in Dhaka [17]. Similarly, an extensively drug resistant *Citrobacter freundii* isolate was identified from a patient returning from India, which was shown to possess nine β-lactamase genes including NDM-1, CTX-M-15, OXA-1, OXA-9, OXA-10, TEM-1, VIM-4 and CMY genes, and an aminoglycoside resistance gene, namely armA [50]. Another study from our neighboring country, Pakistan found out that majority of NDM-1–producing *Enterobacteriaceae* co-produced CTX-M-1 and *armA-*type methylase [51]. These all prove that accumulation of multiple antibiotic resistance genes in the clinical isolates raises an alarm on potential spread of extensively drug-resistant organisms in healthcare settings.

Finally, our data showed many antibiotics with little efficacy against disease-specific pathogens that have acquired resistane genes, especially multiple resistance genes. Not only these antibiotics should be discouraged to use for therapeutic purposes, there should be strict regulation on which antibiotics should be available in the market and which should not be [52, 53]. It is hypothesized that resistant pathogens may shed registance gene(s) without exposure to the antibiotics for a relativey longer period of time and this might be the mechanism for regaining the efficacy by many antibiotics that were in use previously but discontinued due to higher rates of non-susceptilibity.

The study had a number of limitations. We analyzed only ESBL, macrolide and aminoglycoside resistance genes. Antibiotic resistance genes, such as genes for tetracyclines, quinolones and other antibiotics should have been investigated for the isolates analyzed as they were found to be resistant on disk diffusion analysis. The reason behind the genotypic discrepancies for some isolates lies in the fact that there might be the presence of other genes or mutations in the resistance genes which were not investigated in the present study. The isolates that remained undetected with the screening recommendations from CLSI are problematic, since failure to detect ESBL, aminoglycoside and macrolide resistance enzymes can have serious consequences for treatment of the patients [54]. In addition, presence of atypical resistance genes in the organisms should have also been investigated. Furthermore, the study depended on disk diffusion method for preliminary resistance assay. It would have been better to determine the minimal inhibitory concentration (MIC) which could reflect resistance condition more accurately. That is, both MIC and disk diffusion assays might reflect true phenotypes. Moreover, it would offer more clearer information on antibiotic resistance if we could expand investigations by including more hospitals, both within the capital city and the broader territories for active surveillance by increasing the sample size. Now-a-days, whole genome sequencing (rWGS) has been popular for such surveillance studies. However, we could not perform WGS because of resource limitations. This is why follow-up studies are needed indeed to find out whether the resistance genes are plasmid-encoded or not. On top of that, further research and in-depth analysis are required to find out ways and means to solve global antibiotic resistance problem. The research should also focus on how to facilitate resistance gene shedding and inhibit acquisition of new resistance gene.

## Conclusion

In summary, highly pathogenic MDR strains were detected from infected patients in tertiary hospitals of the capital of Bangladesh, which can also contribute to other hospital acquired infections. It is to be noted, however, that these observations reflect the overuse and misuse of antibiotics and underscore the need for urgent steps to arrest the increasing incidence of resistance to the antibiotics in the hospital settings. A tailored antibiotic treatment is in demand to tackle this major issue and regular surveillance should be carried out to monitor the susceptibility of these pathogens and choose appropriate regimens both for prophylaxis and treatment of such hazardous infections.

